# Circular RNAs in the ageing African turquoise killifish

**DOI:** 10.1101/699694

**Authors:** Franziska Metge, Yumi Kim, Jorge Boucas, Christoph Dieterich, Dario Riccardo Valenzano

## Abstract

CircRNAs are a subgroup of RNAs which form a circular molecule. During the splicing process the 3’ splice donor loops back to form a covalent bond with an upstream 5’ splice acceptor instead of the downstream 5’ acceptor. The majority of circRNAs are non-coding splice isoforms of protein coding genes showing tissue and time specific expression. Because circRNAs have no 5’ cap nor 3’ poly-A tail, they degrade slower than their linear host genes. Only few studies were able to show a specific function for circRNAs.

In this work we sequenced 23 samples from three tissues (brain, muscle, and gut) at three (two for gut) time points throughout the life of the naturally short-lived African turquoise killifish (*Nothobranchius furzeri*). We identified 1810 unique circRNAs, half of which are conserved with humans and mice. With this study we provide a comprehensive atlas to the circRNA landscape in the ageing African turquoise killifish, which serves as a novel resource to the circRNA as well as the ageing community.

## 1 Introduction

Circular RNAs (circRNAs) are a special class of RNA generally thought to be non-coding. In contrast to all other spliced RNA classes, circRNAs arise when a 3’ splice donor loops back to form a covalent bond with an upstream 5’ splice acceptor instead of a linear downstream 5’ splice acceptor (see Figure 6 in [Salzman et al., 2012]). Having neither a cap nor a poly-A tail, circRNAs are more stable than linear RNAs [Enuka et al., 2015]. The majority of circRNAs are untranslated splice isoforms of protein coding genes which are called ‘host genes’.

Before 2010, only a few groups explored the circRNA landscape in selected organisms [Hsu and Coca-Prados, 1979, Nigro et al., 1991, Cocquerelle et al., 1992, Capel et al., 1993] but through improved sequencing strategies, circRNAs recently came back into the focus of molecular and computational biologists. Until today, every living organism that has been studied in the context of circRNAs, from yeast and cell cultures to mice and monkeys, express circRNAs [Wang et al., 2014]. Although the exact mechanism of how circRNAs arise remains unknown, circularization of RNA happens co-transcriptionally, depends on the canonical splicing machinery, and uses canonical splice sites [Ashwal-Fluss et al., 2014, Starke et al., 2015]. Intron sequences, intron lengths and transcription elongation rates influence the circularization efficiency [Ivanov et al., 2015, Liang and Wilusz, 2014, Zhang et al., 2016], suggesting specific regulation of circular RNAs in a tissue and time-dependent manner.

Because circRNAs have a longer half life than their linear host genes, they are unlikely to be involved in signaling cascades of fast and dynamic processes but rather could be involved in long-term processes such as differentiation and cell ageing [Enuka et al., 2015]. A study in flies shows a continuous increase in circRNAs in the fly’s nervous system during ageing [Westholm et al., 2014], while studies of embryonic mouse and porcupine brains show a change in circRNA expression during development of the nervous system [You et al., 2015, Venø et al., 2015]. However, due to the lack of appropriate models, there have not been any studies on circRNA expression throughout the whole life span of a vertebrate.

In this study we used the African turquoise killifish as a model organism to study circRNA expression across different tissues during ageing. The African turquoise killifish recently became a valuable model organism for ageing because of its naturally short life span of three to six months while having typical hallmarks of ageing such as neurodegeneration, reduction of locomotion, and loss of pigmentation also present in humans [Harel and Brunet, 2015, Kim et al., 2016, Terzibasi et al., 2007].

In this study we generated an atlas of circRNA expression throughout the life span of the African turquoise killifish. We believe that it provides a resource to researchers studying circRNAs expression during ageing as well as researchers using the African turquoise killifish as a model organism.

## 2 Results and Discussion

To study the circRNA landscape of the ageing African turquoise killifish, RNAseq libraries of three different tissues at three different time points were generated (see Table 1).

**Table 1:** Sample table

| Tissue | Time point | RNA library | Accession |
| --- | --- | --- | --- |
| brain | 3 x 3.5 weeks | 250 BP PE ribo- | GSM3935856, GSM3935857, GSM3935858 |
|  | 3 x 8.5 weeks | 250 BP PE ribo- | GSM3935859, GSM3935860, GSM3935861 |
|  | 3 x 14 weeks | 250 BP PE ribo- | GSM3935862, GSM3935863, GSM3935864 |
| gut | 4 x 6 weeks | 100 BP PE ribo- | SRR5344350, SRR5344349, SRR5344348, SRR5344347 |
|  | 4 x 16 weeks | 100 BP PE ribo- | SRR5344346, SRR5344345, SRR5344344, SRR5344343 |
| muscle | 2 x 3.5 weeks | 250 BP PE ribo- | GSM3935865, GSM3935866 |
|  | 2 x 8.5 weeks | 250 BP PE ribo- | GSM3935867, GSM3935868 |
|  | 2 x 14 weeks | 250 BP PE ribo- | GSM3935869, GSM3935870 |

### More than a quarter of circRNAs are expressed in all tissues

Accounting for all circRNAs found in any sample at the given cutoff, the killifish expresses 1810 different circRNAs across brain, gut, and muscle. A quarter of these are expressed in all three tissues, another quarter of these are expressed only in brain, and six and seven percent are expressed only in muscle and gut respectively. One third of all RNAs are shared among two of the three tissues (Figure 1).

**Figure 1:**
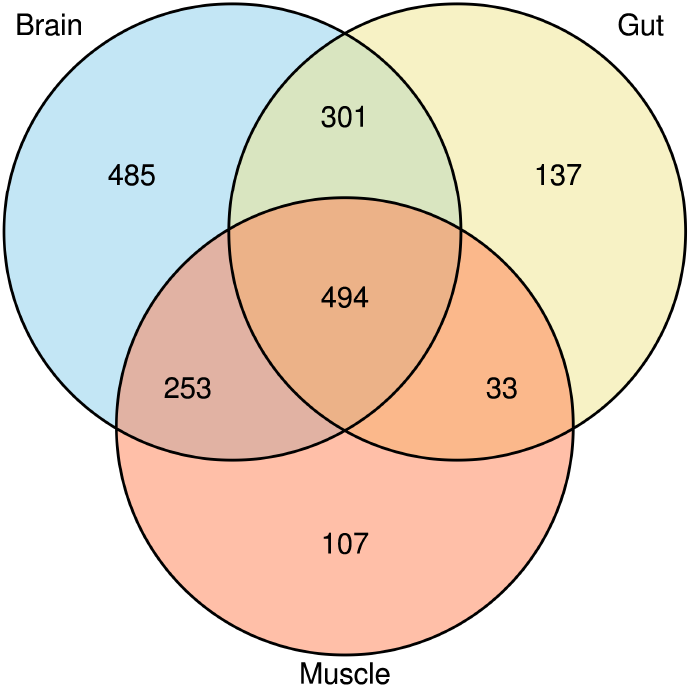
Tissue Overlap. The overlap of detected circRNAs based on their splice coordinates reveals a total of 1810 unique circRNAs across all samples. 494 unique circRNAs are shared among all tissues while 485 unique circRNAs are specific to brain.

### Increased circRNA diversity in the brain

The killifish brain expresses the most diverse amount of circRNAs among the three sequenced tissues (see Figure 2a). This observation supports previous findings by Westholm *et al.* and You *et al.* [Westholm et al., 2014, You et al., 2015]. However, each of these expressed brain circRNAs are on average less abundant than circRNAs in the other tissues (*p*_*brain.vs.muscle*_ = 3.3^−144^, 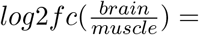 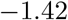; *p*_*brain.vs.gut*_ = 2.7^−8^, 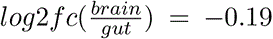, compare Figure 2b). One possibility for this observation could be that circRNAs are highly expressed in one cell type and not expressed in other cell types, thus resulting in a low circRNA/linearRNA ratio. The other possibility could be that circRNAs are lowly expressed throughout all cell types. Considering the amount of circRNAs and the expression level of circRNAs in muscle, the first scenario is more reasonable. The muscle has fewer cell types and fewer host genes expressing circRNAs overall, but at a significantly higher circRNA/linearRNA ratio than the brain. The circRNA landscape of the gut lies in between the brain and the muscle with significantly fewer host genes expressing circRNAs than brain and at a significantly lower ratio than muscle (*p*_*brain.vs.gut*_ = 6.3^−4^, *p*_*gut.vs.muscle*_ = 1.8^−43^, compare Figure 2 a and b).

**Figure 2:**
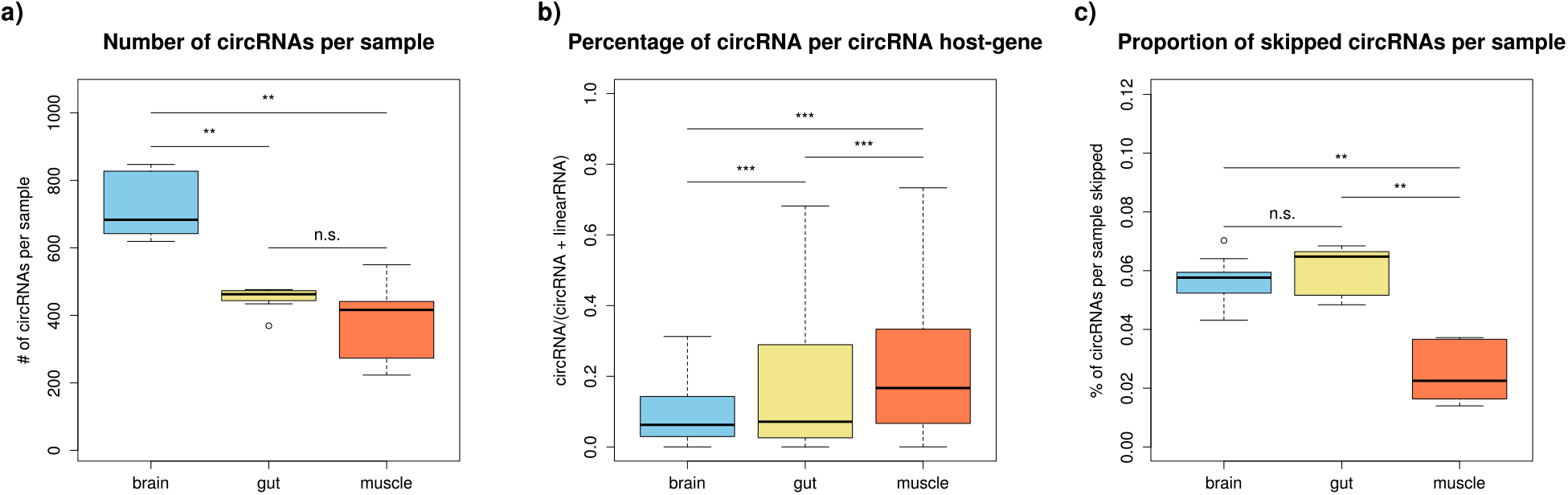
CircRNA landscape. a) Number of different circRNAs shows that the brain exhibits the most diverse circRNA landscape while there is no significant difference between gut and muscle. b) circRNA expression normalized to host gene expression shows that muscle circRNAs have a higher circRNA/host gene ratio than brain and gut. c) Proportion of circRNAs which are being skipped shows that there are significantly fewer circRNAs skipped in muscle than in brain and gut. Wilcoxon Test. * : *p* < 0.01; ** : *p* < 0.001; *** : *p* < 0.0001

Circle skipping are events where there is a linear mRNA isoform excluding exactly the circularized exons [Cheng et al., 2016]. These events could indicate that skipped circRNAs are merely a by-product of the linear splicing. In brain and gut approximately 5% of host-genes exhibit such linear mRNA isoforms, while in muscle only two percent of host-genes express these linear mRNAs skipping circRNA exons (Fig. 2c).

### There are several age dependent circRNAs

The majority of circRNAs peak in abundance during adulthood (Figure 3a,b). In the gut, only two time points were observed revealing five significantly down-regulated and three significantly up-regulated circRNAs in old versus young fish with respect to their host gene (Figure 3c).

**Figure 3:**
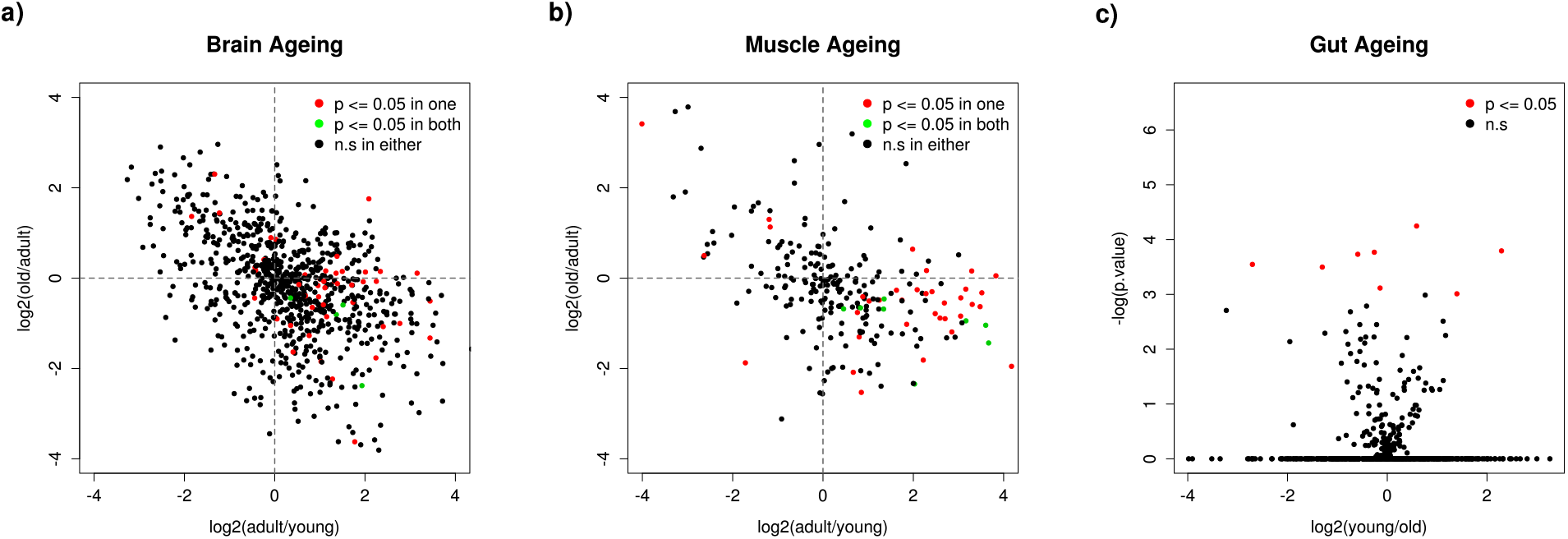
a and b) CircRNA expression during ageing in brain and muscle. CircRNA expression was first normalized by host gene expression. Second, the log2 fold change was calculated to compare the changes from young to adult (x-axis) and adult to old (y-axis). CircRNAs continuously increasing with age are located in the upper right quadrant while circRNAs continuously decreasing with age are located in the lower left quadrant. Figures a and b show that the abundance of the majority of circRNAs peaks in adult killifish. c) circRNA expression in the gut. Young vs. old in relation to p.value shows that only few circRNAs are significantly changed, without any bias to either young or old.

### Gut circRNAs are shorter than brain and muscle circRNAs

CircRNAs expressed in the gut are on average 150-200BP shorter than in brain and muscle (p_*brain*_ = 0.017, *p*_*muscle*_ = 0.0007, see Figure 4). This significant difference can be explained by gut circRNAs having shorter exons as well as fewer exons.

**Figure 4:**
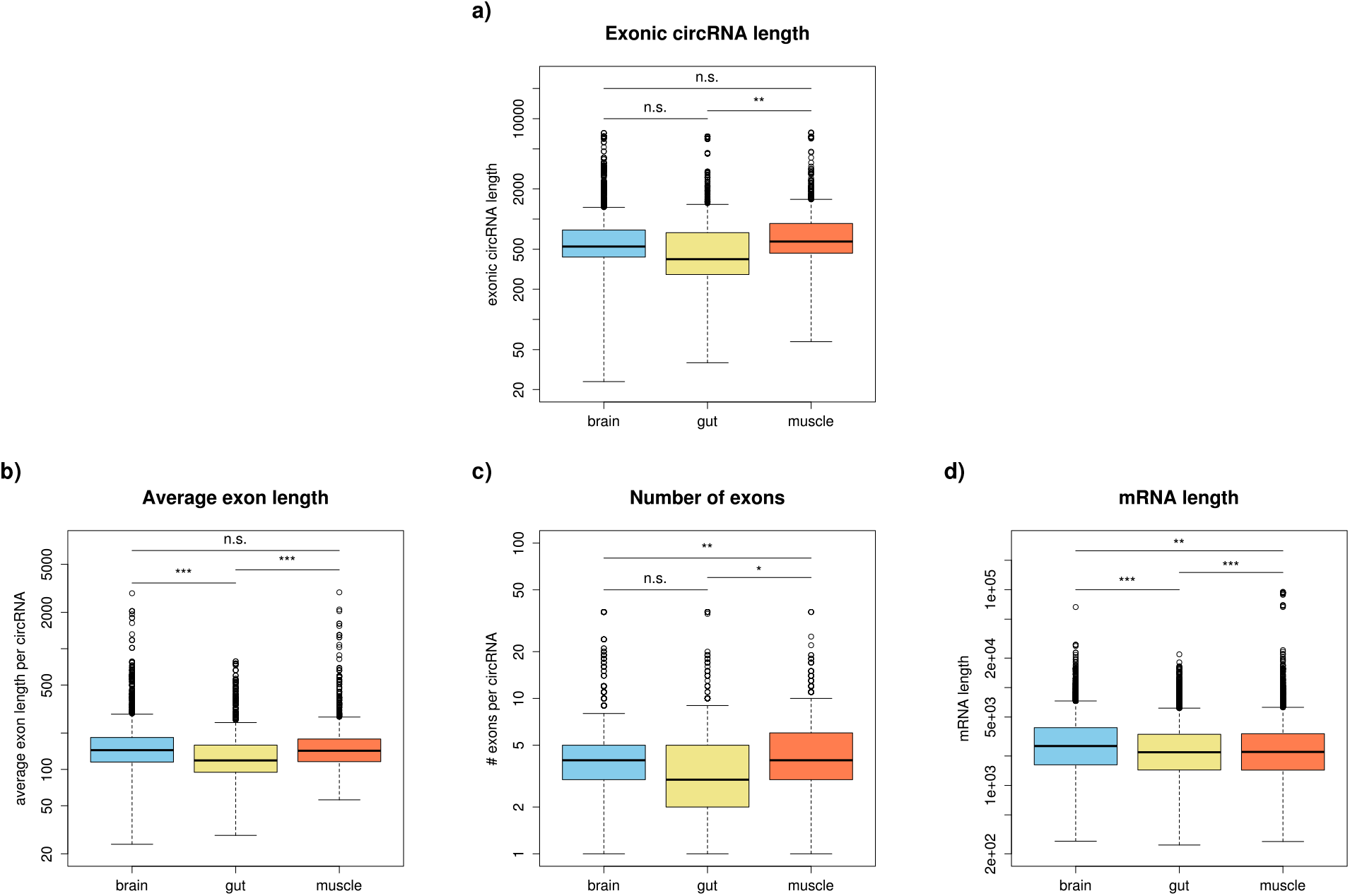
Circle length. a) CircRNA length without introns. Average length: brain = 532 BP, gut = 398 BP, muscle = 597 BP; T-test p-value: brain vs. gut = 0.017, gut vs. muscle = 0.0007, brain .vs. muscle = 0.1. b) Average length per exon over all circRNAs. Average exon length: brain = 144, gut = 119, muscle = 143; T-test p-values: brain vs. gut < 2.2 * 10^−16^, gut vs. muscle = 3.4 * 10^−11^, brain vs. muscle = 0.93. c) Number of exons per circRNA. Average number: brain = 4, gut = 3, muscle = 4; T-test p-values: brain vs. gut = 0.82, gut vs. muscle = 0.002, brain vs. muscle = 0.00015. d) mRNA length without introns. Average length: brain = 2526 BP, gut = 2183 BP, muscle = 2200 BP; T-test p-values: brain vs. gut < 2.2 * 10^−16^, gut vs. muscle = 8.2 * 10^−6^, brain vs. muscle = 0.0004. T-Test. * : *p* < 0.01; ** : *p* < 0.001; *** : *p* < 0.0001

Surprisingly, we find no significant difference in circRNA length between brain and muscle although linear RNAs expressed in the brain are significantly longer than in muscle (p_*circRNA*_ = 0.11, p_*linear*_ = 0.0004).

### More than half of all killifish circRNAs are conserved in other species

We used the reconstructed sequences to align the killifish circRNAs to known circRNAs of *C.elegans*, *D.melanogaster*, *M.musculus*, and *H.sapiens* ^1^. Next we calculated the sequence identity between each killifish circRNA sequence to all other circRNAs from one of the four species. Considering all circRNAs whose sequence is conserved with at least 50% sequence identity, 53% of all killifish circRNAs are conserved in at least one of the four organisms. 48% of killifish circRNAs are conserved in human (867 circRNAs), 26% are conserved in mouse (483 circRNAs), 6% are conserved in fly (109 circRNAs), and 3.5% are conserved in worms (63 circRNAs) (compare Figure 5). Intriguingly, we observe a small proportion of fish circRNAs conserved in mouse but not human.

**Figure 5:**
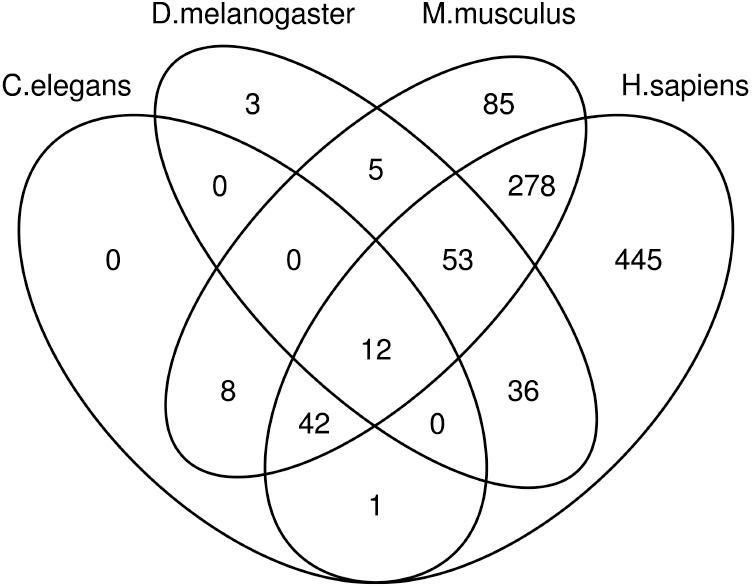
Conservation of killifish circRNA sequences. This figure shows the overlap of conserved killifish circRNAs in other species, e.g. 385 killifish circRNAs are conserved in both human and mouse. However, there are more circRNAs conserved between mouse and human which are not shown here. The largest amount of killifish circRNAs is conserved in humans, followed by mouse, fruit fly, and worm.

## 3 Conclusion

Here we present the circRNA landscape of the ageing African turquoise killifish in brain, gut, and liver. We found a total of 1810 unique circRNAs. While more than a quarter of these are shared among the three tissues, one quarter of these is specifically expressed in brain (Fig. 1). Thus, the brain harbours the most diverse circRNA landscape of all tissues. On the other hand, the circRNA/host-gene ratio is highest in muscle (Fig 2). Furthermore, we could not find an accumulation of circRNAs with age but rather a peak of circRNA expression in adult killifish (Fig 3).

Comparing the exonic circle length revealed that gut circRNAs are significantly shorter and constitute of fewer exons than brain and muscle circRNAs (Fig 4). Blasting the killifish circRNA sequences to known circRNAs downloaded from circbase showed that 53% of killifish circRNAs are conserved across other species with a sequence identity of at least 50% (Fig 5).

In conclusion, by cataloguing the circRNA landscape in the African turquoise killifish, this study provides a resource not only for the circRNA research community but also for the killifish research community.

## 4 Material and Methods

### Fish tissue collection

The short-lived strain of the turquoise killifish (GRZ-AD) is used for this study. The fish were raised as described in [Dodzian et al., 2018]. Brain and muscle tissues were carefully dissected under the microscope. Isolated tissues were briefly rinsed with 1 x phosphate-buffered saline (PBS) and have frozen in liquid nitrogen.

### RNA preparation and sequencing

Biologically independent samples were prepared by pooling five brains or muscles from 3.5-week-old and 8.5-week-old male fish, and four brains or muscles from 14-week-old male fish. Three pools for brain and two pools for muscle were used for RNA isolation by using Trizol (10296028, Invitrogen, USA). RNA quantity and quality were analyzed with an automated electrophoresis system (ExperionTM, BioRad, USA). RNA samples which have over RNA integrity number 8 were sent to Max Planck-Genome-centre Cologne for library preparation and sequencing. Briefly, the sequencing library were prepared after ribosomal RNAs removal using Ribo-zero rRNA removal Kit against human, mouse and rat (Illumina, USA). Sequencing has been done with 2×250 bp paired-end on a HiSeq 2500 (Illumina, USA).

To get a more comprehensive picture of the circRNA landscape previously published gut samples were included [Smith et al., 2017].

### Read trimming

In order to use `STAR` [Dobin et al., 2013] for mapping, reads had to be trimmed to a maximum of 249 base pairs. Additionally reads with more than 5 uncalled bases and reads less than 100 base pairs were removed using `flexbar` [Dodt et al., 2012]:

~~~
flexbar -n 4 -r [sample]_1.fastq.gz -p [sample]_2.fastq.gz -t trimmed/[sample]
~~~

~~~
-f sanger -u 5 -m 100 -k 249 -z GZ
~~~

### Mapping

All RNA-Seq reads were mapped using `STAR` version 2.5.2b to the killifish genome version NFZv2 [Willemsen, 2019] with the following parameters:

~~~
STAR --readFilesCommand zcat --runThreadN 12 --genomeDir [STAR_2.5.2b]
--outSAMtype BAM SortedByCoordinate
--readFilesIn trimmed/[sample]_1.fastq.gz trimmed/[sample]_2.fastq.gz
--outFileNamePrefix star_out/[sample]. --quantMode GeneCounts
--genomeLoad NoSharedMemory --outReadsUnmapped Fastx
--outSJfilterOverhangMin 15 15 15 15 --alignSJoverhangMin 15
--alignSJDBoverhangMin 10 --outFilterMultimapNmax 20 --outFilterScoreMin 1
--outFilterMismatchNmax 999 --outFilterMismatchNoverLmax 0.05
--outFilterMatchNminOverLread 0.7 --alignIntronMin 20 --alignIntronMax 1000000
--alignMatesGapMax 1000000 --chimSegmentMin 15 --chimScoreMin 15
--chimScoreSeparation 10 --chimJunctionOverhangMin 15 --twopassMode Basic
--alignSoftClipAtReferenceEnds No --outSAMattributes NH HI AS nM NM MD jM jI XS
--sjdbGTFfile NFZv2.0.gtf
~~~

Since `STAR` is not able to map read pairs with two chimeric break points we re-aligned all unmapped reads in single end mode using the same parameters as before, suggested in the `DCC` [Cheng et al., 2016] documentation.

### CircRNA detection

CircRNA candidates were detected using `DCC` version 0.4.4 with the NFZv2.0.gtf annotation file. CircRNAs were detected with at least two reads in at least two replicates:

~~~
DCC @sample_sheet -mt1 @mate1 -mt2 @mate2 -D -R NFZv2.0.repeats.sorted.gff
-an NFZv2.0.gtf -B @bamfiles -T 10 -Pi -t tmp/ -F -M -Nr 2 2 -fg -G
-A NFZv2.0.fasta
~~~

### Differential circRNA analysis

Before testing for differentially expressed circRNAs, the list of candidates was filtered to guarantee enough data for useful testing. Only circRNAs obtaining at least three reads in at least three (brain and gut) or two (muscle) samples from one tissue were selected for testing. The beta-binomial test in the `R` package `ibb` [Pham et al., 2010] was used to identify differentially expressed circRNAs. To ensure that the differential expression of circRNAs is not based on an underlying differential expression of host genes, circRNA counts were tested with respect to the sum of linear and circular RNA read counts (see `DCC` for information on counting linear RNA).

### Reconstruction of expressed circle structure

To reconstruct the exact sequence of each circRNA we first extracted all read pairs spanning a back splice junction at least once. Then a list of all linear splicing signals within the circle junction coordinates was generated, sorted and introns were chained together. To create transcripts, exons were inferred in between two introns if they had at least one supporting read. Regions without read support were intersected with annotated exons. Transcripts were saved in `bed` and `bed12` format. `bedtools getfasta` was used to obtain circRNA fasta sequences from these bedfiles. For more information on the reconstruction algorithm see [Metge, 2018]

### Estimating circle length

The length of each circRNA was calculated based on the reconstructed circRNA structures by summing up the block sizes in the `bed12` files. The number of exons was taken from the `bed12` files directly. The average exon length was defined as 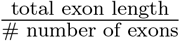 for each circRNA. For Figure 4, each circRNA in each sample was considered.

To compare the circRNA length to linear RNA length we ran `cuffquant` and `cuffnorm` (Cufflinks version 2.2.1. [Trapnell et al., 2010]) using their default parameters. The linear RNA length was extracted from the `isoform.attr_table` and overlapped with the `isoform.fpkm_table` in order to obtain the length of expressed mRNAs only.

### Conseravtion across different species

CircRNA annotations of *H. sapiens* [Salzman et al., 2013, Memczak et al., 2013, Zhang et al., 2013, Jeck et al., 2013], *M. musculus* [Memczak et al., 2013], *D. melanogaster* [Ashwal-Fluss et al., 2014], and *C. elegans* [Memczak et al., 2013] were downloaded from circbase [Glažzar et al., 2014]. Circbase provided files in `bed12` format. These files were supplied `bedtools getfasta` to generate circRNA sequence files. All reconstructed killifish circRNA sequence files were blasted against all reference circRNA sequences using `blastn` (BLAST version 2.2.31+ [Altschul et al., 1990]).

~~~
blastn -gapextend 2 -gapopen 5 -word_size 11 -evalue 10 -penalty -3 -reward 2
-query [killifishFasta] -subject [refFasta] -html -outfmt 0
> [killifishFasta]_[refFasta].blast.out
~~~

Blast results were analyzed for the longest match. A killifish circRNA was considered conserved in a given species if at least 50% of its sequence was matching to any reference circRNA. Figure 5 shows the distribution of conserved killifish circRNAs among the four reference species.

## Footnotes

1 downloaded from http://www.circbase.org/ [Glažzar et al., 2014]

## Notes

https://www.ncbi.nlm.nih.gov/geo/query/acc.cgi?acc=GSE134065

